# Differential neural plasticity of individual fingers revealed by fMRI neurofeedback

**DOI:** 10.1101/2020.03.02.973586

**Authors:** Ethan Oblak, Jarrod Lewis-Peacock, James Sulzer

**Author notes:** These authors contributed equally to this work (Jarrod Lewis-Peacock, James Sulzer).

## Abstract

Previous work has shown that fMRI activity patterns associated with individual fingers can be shifted by temporary impairment of the hand. Here, we investigated whether these neural activity patterns could be modulated endogenously and whether any behavioral changes result from this modulation. We used decoded neurofeedback in healthy individuals to encourage participants to shift the neural activity pattern in sensorimotor cortex of the middle finger towards the index finger, and the ring finger towards the little finger. We first mapped the neural activity patterns for all fingers of the right hand in an fMRI pattern localizer session. Then, in three subsequent neurofeedback sessions, participants were rewarded after middle/ring finger presses according to their activity pattern overlap during each trial. A force-sensitive keyboard was used to ensure that participants were not altering their physical finger coordination patterns. We found evidence that participants could learn to shift the activity pattern of the ring finger but not of the middle finger. Increased variability of these activity patterns during the localizer session was associated with the ability of participants to modulate them using neurofeedback. Participants also showed an increased preference for the ring finger but not for the middle finger in a post-neurofeedback motor task. Our results show that neural activity and behaviors associated with the ring finger are more readily modulated than those associated with the middle finger. These results have broader implications for rehabilitation of individual finger movements, which may be limited or enhanced by individual finger plasticity after neurological injury.

## 1 Introduction

Movements of individual fingers have distinct neural activity patterns that can be measured using multi-voxel pattern analysis (MVPA) of fMRI data from sensorimotor cortex (Diedrichsen et al., 2012; Ejaz et al., 2015; Kolasinski et al., 2016a). In animal models, the underlying neural activity maps in sensorimotor cortex have long been known to adapt dynamically with experience and learning (Buonomano and Merzenich, 1998; Feldman and Brecht, 2005). More recently, it was found that neural finger representations in humans will adapt and shift when two of the fingers are glued together for a short period of time (Kolasinski et al., 2016b). Furthermore, this temporary adaptation of cortical maps was correlated with behavioral deficits recorded after the fingers were unglued.

Instead of physically intervening at the hand (e.g. by gluing fingers together) and measuring the associated cortical changes, our approach is to directly induce cortical remapping and observe whether any changes in motor behavior or tactile sensation of the fingers occur. Current neural stimulation techniques such as transcranial magnetic stimulation (TMS) and transcranial direct-current stimulation (tDCS) operate on the scale of centimeters, and are therefore unable to target the millimeter-scale neural activity patterns associated with individual finger movements in humans (Walsh and Cowey, 2000; Brunoni et al., 2012). However, a recent technique known as decoded neurofeedback is able to target activity patterns directly, for example to modulate the fMRI patterns associated with different visual orientations in primary visual cortex (Shibata et al., 2011). Critically, the modulation of these patterns correlated with changes in visual perception, establishing a link between direct modulation of neural activity patterns and their associated behavior. This technique has since been used to investigate neural representations such as fear conditioning (Koizumi et al., 2016; Taschereau-Dumouchel et al., 2018) and confidence judgments (Cortese et al., 2016), but has yet to be used to target motor behaviors or tactile sensation.

Therefore, we wanted to investigate whether decoded neurofeedback could be applied to neural maps in sensorimotor cortex and whether this would result in behavior changes similar to those found in physical manipulations such as gluing or repeated stimulation of individual fingers. Instead of physically attaching the fingers together, we used a form of decoded neurofeedback known as associative decoded neurofeedback (Amano et al., 2016). This technique was used by Amano et al. (2016) to associate a neutral visual stimulus (grey orientation grating) with another visual stimulus (colored orientation grating). In this study, we used associative decoded neurofeedback to associate presses of one finger (either the middle or ring finger) with the outwardly adjacent finger (index or little finger). This manipulation was intended to match the neural changes found by Kolasinski et al. (2016b), in which the representations of the middle and ring finger were shifted apart, and the representations of the ring and little finger moved closer together. We hypothesized that individual finger representations and their associated behaviors would be altered after neurofeedback training.

The complete experiment consisted of four fMRI sessions: an fMRI finger pattern localizer session followed by three neurofeedback sessions (Fig 1A). During the pattern localizer, participants pressed each of the fingers an equal number of times, whereas during neurofeedback participants pressed only the middle and ring fingers. Pressing behavior was controlled using a custom force-sensitive keyboard to ensure participants exerted the same amount of force and didn’t alter their finger coordination patterns during each trial (Fig 1B). Motor behavior was assessed before and after each fMRI session using a rapid reaction time task (Fig 1C). This task captured behavioral changes resulting from any short-term plasticity associated with the neurofeedback manipulation as well as any changes due to repeated presses of only the middle and ring fingers over the course of each neurofeedback session. Somatosensory perception was assessed before and after the full experiment using a tactile temporal order judgment task in which adjacent finger pairs were stimulated in quick succession (Fig 1D). This assessment determined if any changes in tactile discrimination ability occurred due to neurofeedback. Overall, this approach offers insight into how changes in neuroplasticity can alter sensorimotor behavior, with implications for recovery following neurological injury such as stroke.

**Figure 1:**
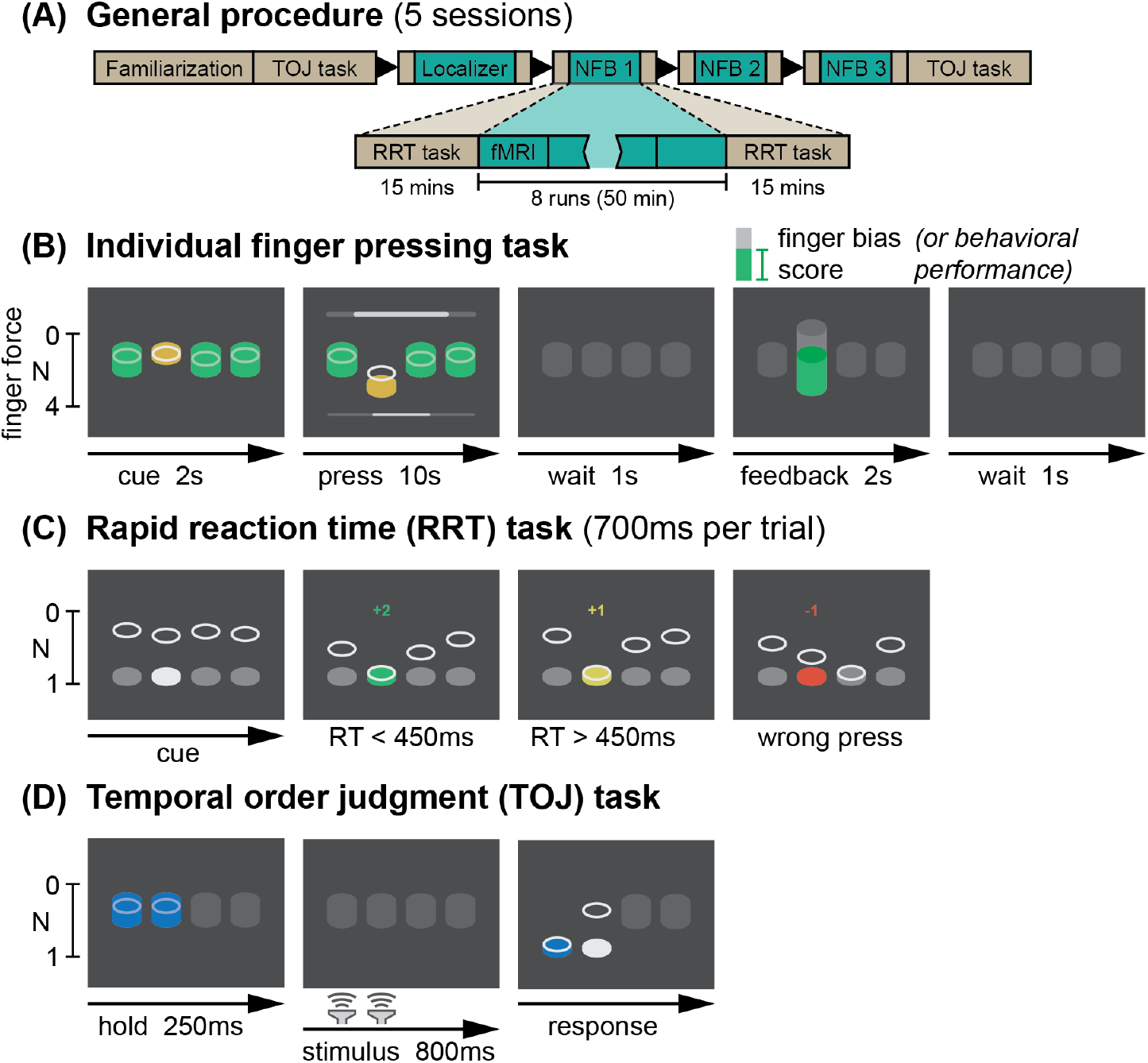
Experimental design. **(A)** The 5-session experiment consisted of behavioral familiarization, a finger pattern localizer fMRI session, and 3 neurofeedback sessions. Each fMRI session included behavioral pre- and post-tests. **(B)** An individual finger pressing task was used as the basis for the localizer and neurofeedback sessions. Participants were required to make individual presses with one of 4 fingers (index, middle, ring, or little) while maintaining constant pressure on all other keys. At the end of each trial, feedback was presented related to their motor behavior (localizer session) or their ability to bias the fMRI patterns related to finger presses (neurofeedback sessions). **(C)** A rapid reaction time (RRT) task was used to assess motor confusion before and after each fMRI session. Participants were encouraged to make rapid presses (reaction time below 450ms) through a point system. **(D)** A temporal order judgment (TOJ) task was used to assess the hand representation of participants before and after the entire neurofeedback protocol. During a 800ms stimulus blank period, a brief vibrotactile stimulus was delivered to 2 adjacent fingers in rapid succession. Participants then judged which of the two stimuli happened first.

## 2 Results

### 2.1 Differential modulation of neural patterns related to individual fingers

Presses of each of the ring and middle fingers were associated with a ‘bias score’ intended to shift their representations based on real-time decoded patterns of fMRI activity. After each 10-sec trial of pressing, this score was displayed to participants as a feedback thermometer (Fig 1B). For the middle finger, the score was higher when the real-time decoded pattern appeared more similar to the index finger, and lower when the decoded pattern was more similar to the ring finger. For the ring finger, the score increased with little finger pattern similarity and decreased with middle finger pattern similarity. Participants received monetary reward related to their ability to increase the bias scores for each finger, up to a total of $30 per session.

Participants were unable to reliably increase the middle finger bias score above the baseline level from the pattern localizer (t_(9)_=−0.22, p=0.83) (Fig 2A, left column). However, participants were able to increase the ring finger bias score above baseline: mean bias scores for the ring finger were significantly greater during neurofeedback compared to the pattern localizer session (t_(9)_=2.63, p=0.027) (Fig 2A, right column). A linear mixed effects model of bias score with participant as a random effect confirmed a significant interaction between session (localizer or neurofeedback) and finger (middle or ring) (F_(1,27)_=5.15, p=0.031). A direct comparison between bias scores also showed significantly higher scores for the ring finger compared to the middle finger during neurofeedback (t_(9)_=2.47, p=0.035).

**Figure 2:**
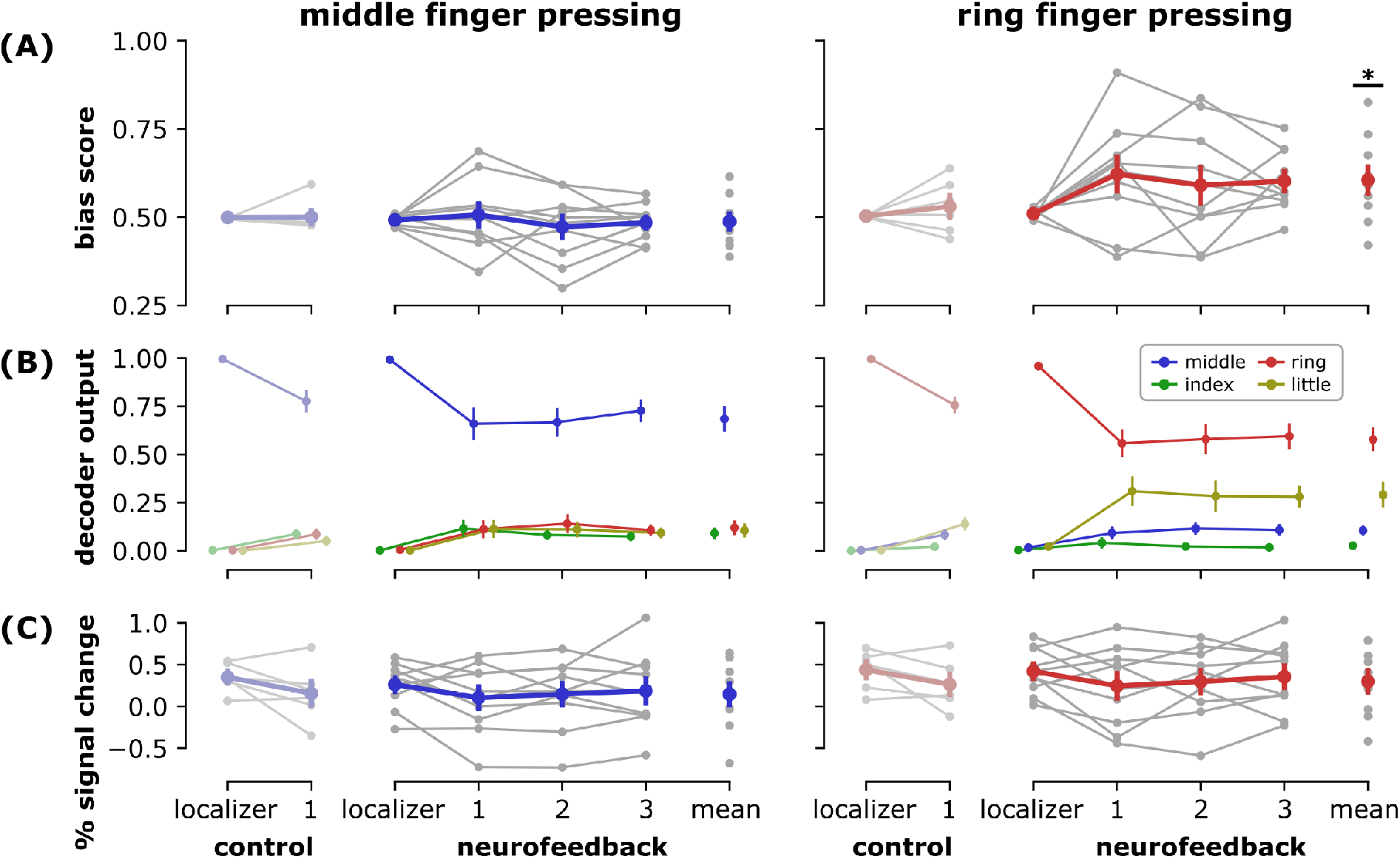
Individual finger pattern bias modulation. **(A)** Mean bias scores by session for presses of middle finger (blue, left column) and ring finger (red, right column). See Equation 1 in Methods for bias score calculation details. **(B)** Mean decoder outputs by session. During middle finger presses, bias scores encouraged participants to increase the index finger decoder output (green) relative to the ring finger output (red). During ring finger presses, bias scores encouraged participants to increase the little finger output (yellow) relative to the middle finger output (blue). **(C)** Mean univariate activations by session, calculated as the mean percent signal change within the voxels selected by each decoder. Univariate activations were found to be unrelated to changes in bias score. Control data were acquired from participants (N=6) that performed two sessions of finger pressing but did not receive neurofeedback in the second session (Oblak et al., 2019). Neurofeedback data (N=10) are shown by session (localizer: pattern localizer; 1-3: neurofeedback sessions; mean: average over all neurofeedback sessions). All data (control and neurofeedback) are shown relative to a decoder trained on each participant’s localizer session. Grey dots indicate means for each participant, colored dots indicate means across the entire group. Errors bars indicate standard error of the mean (SEM). Statistical differences relative to the pattern localizer sessions are indicated at p<0.05 (*).

### 2.2 Changes in univariate activations did not inflate bias scores

Extensive practice of a motor task can result in increases or decreases in univariate neural activation in primary sensorimotor cortex (Steele and Penhune, 2010; Dayan and Cohen, 2011; Hardwick et al., 2013). Changes in univariate activations could potentially drive bias scores, for example if the decoder weights are biased towards one finger or if changes in signal to noise ratio (SNR) alter the decoder’s confidence. We therefore investigated whether the univariate signal (percent signal change; PSC) within the voxels selected by the decoder in each participant were related to their calculated bias scores. We analyzed this relationship in neurofeedback participants as well as in a control dataset where participants performed individual finger pressing in two subsequent sessions but did not receive neurofeedback (Oblak et al., 2019).

We found no significant relationship between mean univariate activation and finger bias score in the control dataset for either the middle finger (linear mixed effects model with participant as random effect; general linear hypothesis test; p=0.705) or ring finger (p=0.664). In the neurofeedback group, this relationship was trending for the middle finger (p=0.072) but not for the middle finger (p=0.334). For the middle finger, the model estimate for the relationship between PSC and bias score was negative for both the control (*β*=−0.022) and neurofeedback (*β*=−0.068) datasets, implying that decreased univariate activity during middle finger presses could increase middle finger bias scores. For the ring finger, this model estimate was positive for both control (*β*=0.041) and neurofeedback (*β*=0.073) datasets, implying that increased univariate activity for the ring finger could increase ring finger bias scores. However, the mean univariate activation of the ring finger during neurofeedback was not increased relative to the pattern localizer session (t_(9)_=−1.45, p=0.18), and the negative estimate of PSC (*β*=−0.12) was in the opposite direction of the potential relationship between PSC and bias score. Therefore, we are confident that epiphenomenal changes in univariate activation did not contribute to apparent neuromodulation success during ring finger presses.

### 2.3 Baseline variability predicts neuromodulation ability

When bias scores were taken from the pattern localizer session for all participants, mean bias scores were similar for the middle (0.494) and ring (0.507) fingers, but variability (standard deviation) was on average larger for the ring (SD=0.038) than the middle (SD=0.027) (Fig 3A). This variability was not reliably different for each finger when compared within subject (t_(9)_=1.67, p=0.13), but did correlate with individual participants’ abilities to increase each finger’s bias score (Spearman’s rho=0.39, p=0.036; see Methods for bootstrapping details) (Fig. 3B).

**Figure 3:**
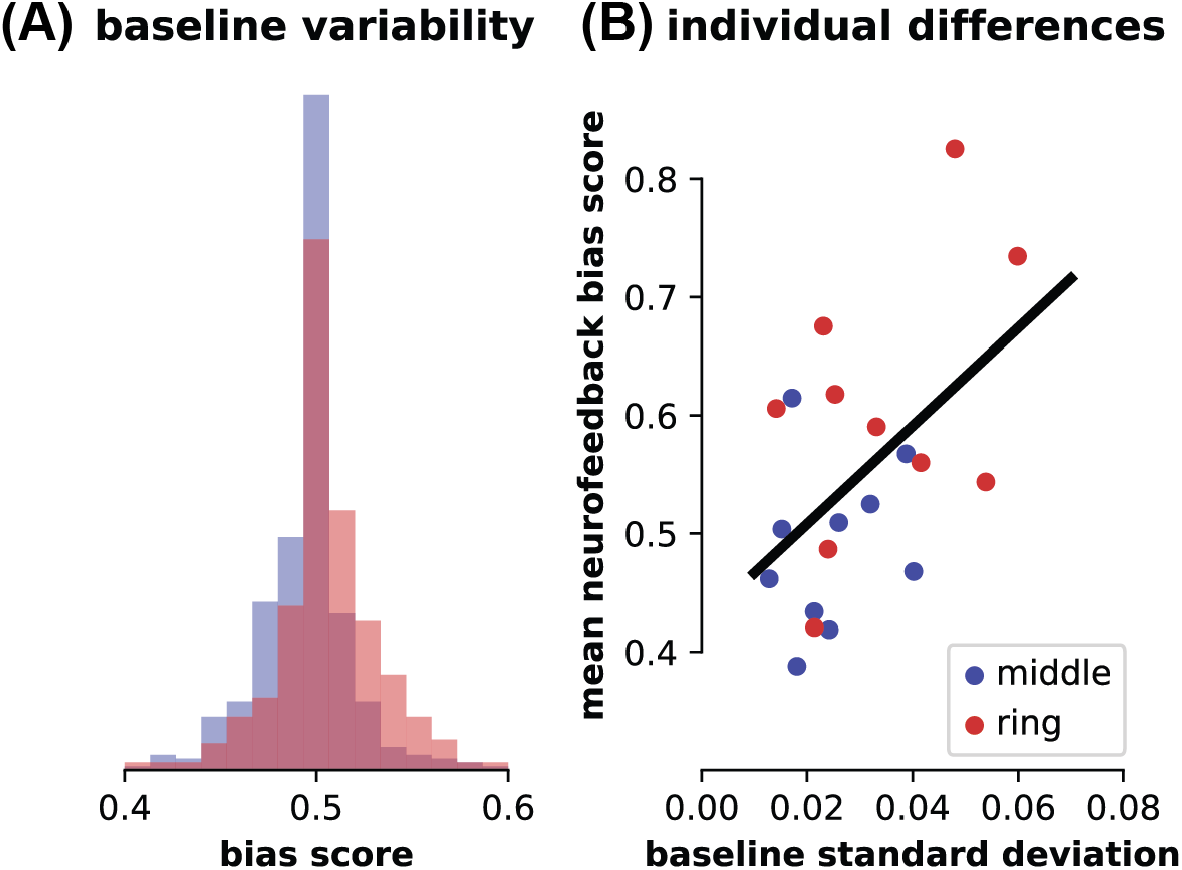
Baseline variability of individual finger patterns. **(A)** Distributions of bias scores for middle (blue) and ring (red) fingers for all participants during the baseline pattern localizer session. **(B)** Relationship between individual participant bias pattern variability during the pattern localizer session (standard deviation) and mean bias score achieved during neurofeedback sessions for each finger. Best-fit line shown for reference; see Methods for statistics.

### 2.4 Neuromodulation was not driven by finger coordination

We used a custom force-sensitive keyboard to ensure that no overt movements were performed with the non-target fingers. It was nevertheless possible that participants could learn to subtly shift their finger coordination to increase the bias scores. We therefore examined whether bias scores were related to finger coordination. We hypothesized that increased movement of the rewarded finger (press middle/reward index; press ring/reward little) would increase bias scores, and increased movement of the punished finger (press middle/punish ring; press ring/punish middle) would decrease bias scores. No such relationship was found for each participant on average (Fig 4, upper panels). Mean coordination patterns with the adjacent fingers in each participant did not explain each participant’s mean neurofeedback scores for either the middle finger (linear model without random effects; general linear hypothesis test; index fixed effect: p=0.838; ring fixed effect: p=0.428) or the ring finger (middle: p=0.986; little: p=0.446).

**Figure 4:**
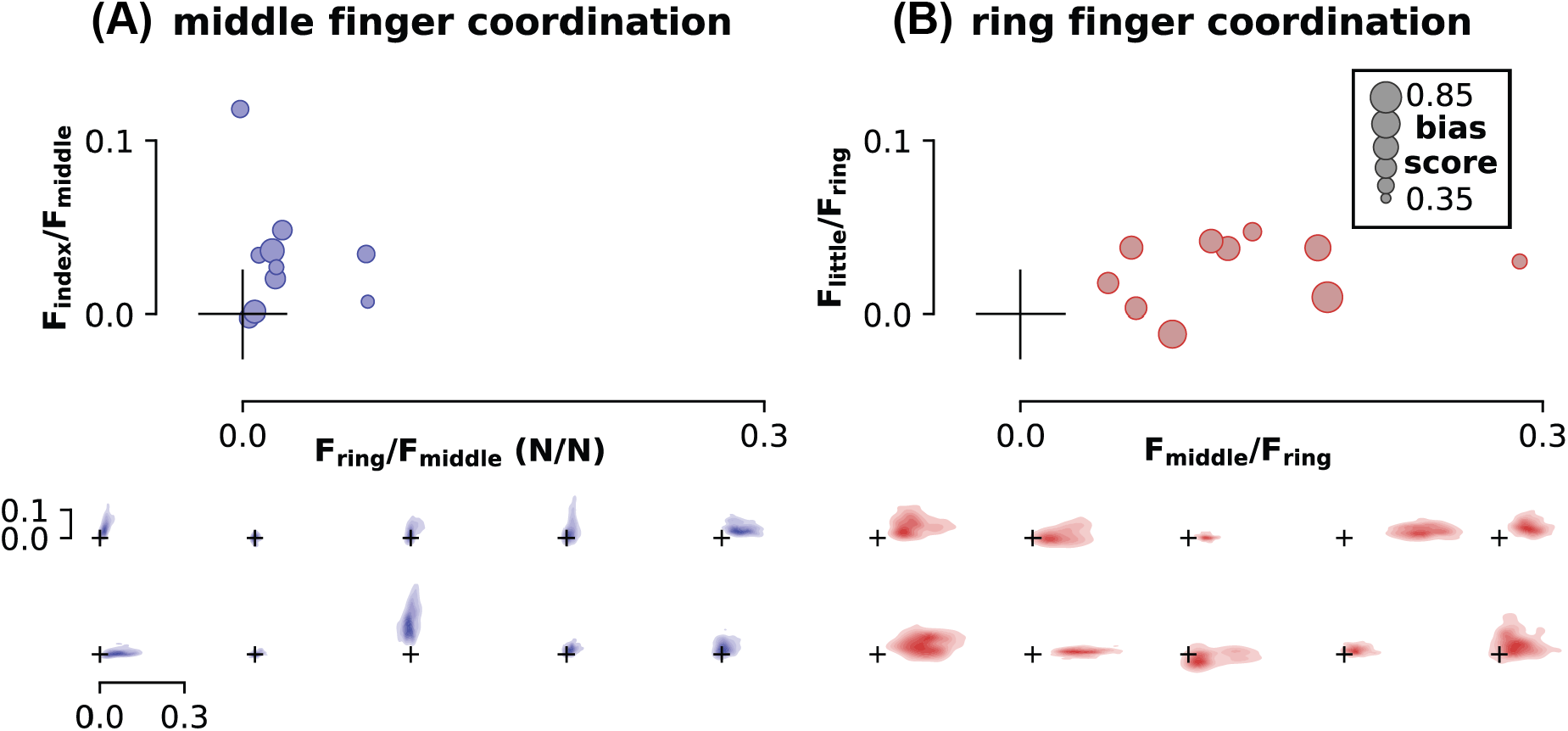
Finger coordination patterns. **(A)** Coordination of middle finger presses with the rewarded finger (index, y-axis) and punished finger (ring, x-axis). **(B)** Coordination of ring finger presses with the rewarded finger (little, y-axis) and punished finger (middle, x-axis). Mean coordination values for each participant (top panel, N=10) are shown as a ratio of the force of the rewarded or punished finger to the pressed finger (N/N). The size of each marker indicates the mean bias score achieved by each participant, with larger circles indicating a greater bias score. These plots illustrate that coupling between adjacent fingers did not correlate with bias scores; if this were true, we would expect larger circles towards the top of each plot (rewarded finger coupling increasing on the y-axis) and smaller circles towards the right of each plot (punished finger coupling increasing on the x-axis). Individual coordination heatmaps for each participant are also shown in the lower panel. Each heatmap in the lower panel corresponds to one circular marker in the upper panel, with the size and position of the origin marker cross serving as a scale reference. See Methods for details of coordination pattern calculation and heatmap generation.

While average coordination patterns for each participant did not predict their mean bias scores, it is possible that within-subject coordination pattern variability could explain bias scores on a trial-by-trial basis. Individuals demonstrated coordination patterns with unique distributions (Fig 4, lower panels). However, we found no relationship between trial-by-trial coordination patterns and bias scores in either the middle finger (linear mixed effects model with participant as random effect; general linear hypothesis test; index fixed effect: p=0.11; ring fixed effect: p=1.00) or ring finger (middle: p=0.924; little: p=0.663).

### 2.5 Participants were unaware of the neurofeedback manipulation

Participants reported their subjective strategies for each finger at the end of each neurofeedback session. Only three of ten participants reported motor-related strategies, including ‘pressing slowly and accurately’, ‘focusing only on the target finger’, and ‘trying not to push both middle and ring fingers’. The remaining seven participants reported various strategies such as ‘thinking about math problems and counting’, ‘relaxing and looking at the top of the thermometer’, and ‘thinking positive thoughts’.

After the final neurofeedback session, participants also reported what they thought the height of the feedback thermometer was related to. Four of ten reported ‘I don’t know’ or ‘no idea’. Only two participants reported different strategies for each finger (first participant: ‘eye focus’ for the middle finger, ‘ring finger pressure’ for the ring finger; second participant: ‘heightened emotions’ for the middle finger, ‘confidence and focus’ for the ring finger). The remaining four participants reported various guesses for the thermometer’s height, with the same guess being reported for both fingers: (1) ‘positive thoughts’, (2) ‘creative thoughts while accurately performing motor task’, (3) ‘ability to isolate finger movements’, and (4) ‘maintaining focus while engaging in higher order logical thinking’.

Finally, it was revealed to participants that increases in the height of the thermometer for each finger was related to one of the non-target fingers, and they were asked to guess which of the three that was. For the middle finger target, three participants guessed the index finger, three guessed the ring finger, and four guessed the little finger. For the ring finger, six participants guessed the index finger, three guessed the middle finger, and one guessed the little finger.

Overall, these subjective results suggest that participants were unaware of the purpose of the neurofeedback manipulation. A minority of participants reported motor-related strategies, which were mostly related to performing the ongoing task accurately (i.e. pressing the target finger independently). Although one participant did report ‘trying not to push both middle and ring fingers’ after one neurofeedback session, this was not reiterated during the final questionnaire regarding the height of the thermometer. Even when it was finally revealed that the neurofeedback reward was related to non-target fingers, participants did not show a bias towards the rewarded fingers when asked to choose between non-target fingers.

### 2.6 Differential modulation of finger preference

Motor behavior was assessed before and after each fMRI session in a rapid reaction time task. The pressing order in this task was randomized to have an equal number of presses of each finger and an equal number of transitions between each possible two-finger combination. We considered two factors that would influence participants’ pressing behavior: the neurofeedback manipulation as well as the fact that participants would only be pressing the middle and ring fingers (and not the index or little fingers) throughout each fMRI neurofeedback session.

To assess the effect of motor repetition, we measured finger preference as the proportion of total presses by each finger during each testing session (Fig 5A). We hypothesized that participants would have increased preference for the middle and ring fingers after each neurofeedback session due to motor repetition. Consistent with this hypothesis, we found that after the first neurofeedback session the preference for the ring finger significantly increased (t_(9)_=3.82, p=0.004) and preference for the little finger significantly decreased (t_(9)_=−3.50, p=0.007). Preference for the index finger was trending towards a decrease (t_(9)_=−1.87, p=0.094), but contrary to our hypothesis, preference for the middle finger did not significantly increase (t_(9)_=1.56, p=0.15). After the second neurofeedback session, preference for the little finger was trending towards a decrease (t_(9)_=−2.19, p=0.056) but no other near-significant results were found (all other p>0.1). After the third neurofeedback session, preferences were unchanged (all p>0.1).

**Figure 5:**
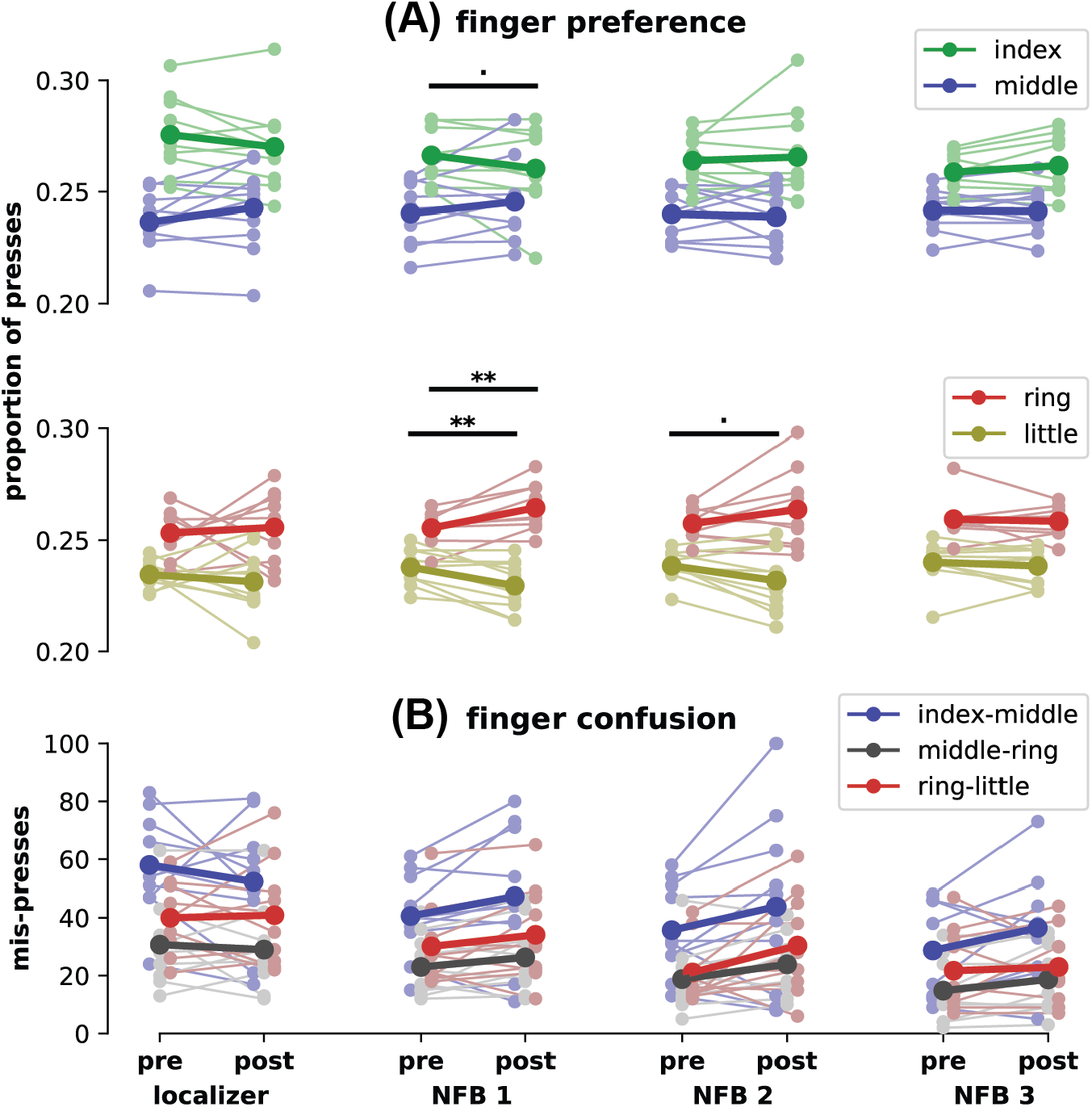
Finger pressing behavior before and after each fMRI scan. **(A)** Finger preference in pre and post-fMRI rapid reaction time tests, as a proportion of the total number of presses. Faded colors indicate each participant’s performance, solid colors indicate means across the group. Within-finger statistical differences from pre to post-test are shown at p<0.1 (.) and p<0.01 (**). **(B)** Confusion between fingers in pre and post-fMRI rapid reaction time tests. Mis-presses are assigned to a finger pair when the target finger was one of the fingers of the pair, but the other finger in the pair was pressed instead.

We also assessed the relative change in finger preference by contrasting the change in preference of adjacent fingers, on a session-by-session basis. For the localizer session and the third neurofeedback session, there were no significant differences in the change in preference between any adjacent fingers (general linear hypothesis test, all adjacent finger contrasts p>0.1). After the first neurofeedback session, the increase in ring finger preference was significant in contrast to the decrease of the little finger (p<0.001). The increase in middle finger preference was also significant in contrast to the index finger (p=0.016). There was no significant difference between the change in ring and middle finger preferences (p=0.63). After the second neurofeedback session, the increase in ring finger preference was still significant relative to the little finger (p=0.049), but there was no significant difference between index and middle (p=0.91) or middle and ring (p=0.38) preference changes.

Both the absolute and relative changes in finger preference were most noticeable after the first neurofeedback session, barely present after the second, and completely gone after the third. By the end of the third neurofeedback session post-test, participants had approximately 120 minutes of exposure to the rapid reaction time task, and their mean pressing accuracy had increased from 81.4% (localizer session, pre-test) to 92.6% (neurofeedback session, post-test). Therefore, it is likely that overtraining on the rapid reaction time task led to decreased sensitivity to finger preference changes during later neurofeedback sessions.

To assess the neurofeedback manipulation, we measured motor confusion between finger pairs using the number of mis-presses within each finger pair. A mis-press was defined as a press that was supposed to be with one finger of the pair, but was instead performed with the other finger of the pair. We hypothesized that the neurofeedback manipulation would result in increased confusion (mis-presses) within index-middle and ring-little finger combinations, and decreased confusion within the middle-ring finger pair, as was found by Kolasinski et al. (2016b). Despite the changes in finger preference found earlier, there were no significant differences between the change in mis-presses in any finger pairs (Tukey’s HSD, all sessions, all finger pair contrasts p>0.1) (Fig 5B). We also assessed somatosensory ability before and after the entire experiment using the just noticeable difference (JND) in tactile discrimination within each finger pair. There were no significant differences between the tactile performance changes in any finger pairs (Tukey’s HSD, all finger pair contrasts p>0.1). These results show that while motor repetition may have caused changes in finger behavior, there is no evidence that increasing pattern overlap through neurofeedback was able to increase confusion between fingers in the motor or sensory domains.

## 3 Discussion

In this study both neural activity and behavior associated with the ring finger were manipulated, while the same was not possible for the middle finger. Participants implicitly learned to shift the neural representation of the ring finger toward that of the little finger, but on average were unable to shift the representation of the middle finger toward the index finger. Moreover, the baseline variability of participants’ individual finger representations were predictive of their ability to bias each finger’s representation. After the first neurofeedback session, participants also showed increased preference for the ring finger and decreased preference for the little finger, although no such preference shifts occurred with the middle or index fingers. These results imply greater plasticity for the ring finger than the middle finger, which is supported by the natural variability of these fingers’ neural representations.

Our results showing differential neural plasticity are complementary to those found by Kolasinski et al. (2016b). Although they physically glued the index and middle fingers together and hypothesized plasticity changes within those fingers, they only found evidence of shifts in the representation of the ring finger, with the strongest behavioral changes also found within the ring and little fingers. These results also agree with animal experiments in which electrophysiologically-recorded signals were easier to control when they aligned with more naturally varying neural activity patterns, rather than along an unnatural manifold (Sadtler et al., 2014). In decoded neurofeedback, there is also within-subject evidence that certain patterns are easier to regulate than others. In a crossover design, Cortese et al. (2017) found that while participants could learn to both up and down-regulate patterns related to confidence judgments, upregulation of these patterns was easier and had a stronger effect on subsequent learning of downregulation compared to the reversed condition. The relationship discovered between neural pattern variability and learning ability supports our previous findings in which pattern localizer data was able to predict participants’ abilities in a subsequent decoded neurofeedback task (Oblak et al., 2019). By examining baseline variability of patterns, future studies may be able to determine the feasibility of controlling specific neural patterns.

While decoded neurofeedback has targeted behaviors such as visual perception, confidence ratings, and fear responses, this work represents the first decoded neurofeedback study to target motor behaviors (for comprehensive overview, see Watanabe et al. (2017)). Furthermore, it is the first fMRI neurofeedback study to target individual fingers, rather than modulating overall sensorimotor cortex activity (Bray et al., 2007; Blefari et al., 2015). These results also have implications for rehabilitation of individual finger movements after stroke. The strength and coordination of individual finger movements are impaired after stroke, recovering significantly through the first year post-stroke but often leaving chronic deficits (Xu et al., 2017). Recent work has suggested that this recovery and subsequent chronic impairment can be tracked through MVPA of individual finger representations (Xu et al., 2015), although the to-date published results of the fMRI data from this study report only average region-of-interest analyses (Ejaz et al., 2018). Although advanced robotic techniques exist for improving finger individuation (Thielbar et al., 2014; Taheri et al., 2014), neurofeedback enables targeted neuroplasticity, and with it, the potential ability to avoid maladaptive neuroplasticity. This approach has a long history of use in altering spinal reflexes in rats (Chen and Wolpaw, 1995), non-human primates (Wolpaw, 1987), and humans (Thompson et al., 2009). More recently, it been used with success to improve gait in spinal cord injury patients without requiring conscious effort (Thompson et al., 2013), showing promise for rehabilitation. As our technique shows the ability for participants to gain control over the neural activity patterns related to individual fingers, we suggest that we may be able to improve individual finger coordination after stroke through the use of decoded neurofeedback.

A major limitation of this study is that there was no observable change in the targeted finger behavior that could be conclusively linked to the neurofeedback manipulation. This is not unlike operant conditioning of spinal reflexes: Thompson et al. (2009) did not find any changes in gait due to conditioning in healthy individuals, and it was only later after studying patients in which significant gait improvements were found (Thompson et al., 2013). This paradoxical improvement is explained through a negotiated equilibrium model: when plasticity that produces a new behavior also improves a previous behavior, overall plasticity is increased, leading to greater changes in patient populations (Wolpaw, 2018). Therefore, it may be necessary to perform decoded neurofeedback training on patients with significant deficits to observe any changes in finger behavior. Furthermore, a major behavioral confound in our study (pressing of only two of the fingers during neurofeedback training) may have masked any small effects of decoded neurofeedback. An ideal experiment would involve training of all four fingers; however, this would involve learning twice as many patterns as the present experiment. Not only would this double the duration of the experiment, it is also unknown whether learning four different neurofeedback targets is feasible within a single experimental group. As this is the first study of its kind, we compromised between too little variation (e.g. only training one finger and losing the within-subject control of two different fingers) and presenting participants with too difficult of a task (the confusion associated with training all four fingers). This allowed us to have a within-subject control (training of 2 different fingers) without presenting a potentially impossible task to participants. As this study shows promise towards controlling multiple patterns, future work should attempt to include all possible fingers. This could be achieved either through careful consideration of the training schedule to ensure participants can learn to control the multitude of patterns, or by including multiple groups, with different finger pairs selected for each group. We also found no changes in sensory perception, which may be due to the neurofeedback task being primarily a motor task. Future studies could include vibrotactile stimulation during the neurofeedback phase to enhance changes in sensory behavior.

Overall, the first attempt at decoded neurofeedback in sensorimotor cortex demonstrated both promise and limitations for the technique. Our results show that certain patterns and behaviors may be difficult to manipulate in certain fingers or in individuals with decreased sensorimotor cortex plasticity. However, our results were able to support existing literature and provide a unique window into the plasticity and representations of individual finger movements in humans. Furthermore, we present a unique technique that may be able to enhance rehabilitation of individual fingers without requiring physical intervention at the limb or knowledge of the manipulation from patients.

## 4 Methods

### 4.1 Participants

Fourteen healthy participants were recruited for the experiment in accordance with the University of Texas at Austin Institutional Review Board. Informed consent was obtained from all participants. 1 participant became uncomfortable during the first fMRI session and did not continue in the experiment. 2 participants completed the first fMRI session but elected not to continue in the experiment. 1 additional participant completed the familiarization session outside of the MRI scanner but did not continue the experiment due to illness. Thus, 10 participants completed the experiment and remained in the final analysis (4 female, average age 25.5 years, SD=5.2). This sample size is in line with previous studies of finger plasticity (Pleger et al., 2003; Kolasinski et al., 2016b). As neurofeedback is a new approach to studying finger plasticity, this study aimed to test the viability of this technique. Future studies will be able provide stronger conclusions with an increased number of participants.

### 4.2 Apparatus

Finger presses were recorded from the index, middle, ring, and little fingers of the right hand using a custom force-sensitive keyboard at 120Hz. For tests outside of the MRI scanner, the keyboard (force sensor: TE FS20) was placed on a desk, and visual instructions and feedback were presented on a laptop screen using Python and Pygame. For tests inside the MRI scanner, the keyboard (force sensor: Honeywell FSS015) was affixed to a wooden board, which lay on the participant’s lap, and visual instructions and feedback were provided through a back-projection screen. For vibrotactile stimuli, two vibrotactile stimulators (Soberton E-12041808) were placed on top of the force-sensitive keyboard.

### 4.3 Imaging parameters

Participants were scanned in a Siemens Skyra 3T scanner with a 32-channel head coil. For all fMRI sessions, the same EPI sequence was used (TR=2s; 36 slices; in-plane resolution 2.3×2.3mm; 100×100 matrix size; 2.15mm slice thickness; 0.15mm slice gap; 2x multiband factor). After auto-alignment to the AC-PC plane, a manual adjustment was performed to ensure coverage of the motor cortex. The same manual adjustment parameters were applied to the subsequent neurofeedback sessions. A high-resolution anatomical scan (MEMPRAGE, 1mm isotropic voxels) was also acquired during the first localizer session to identify the primary sensorimotor cortex (M1+S1) using Freesurfer. Subsequent real-time fMRI scans were rigidly aligned to this template using AFNI’s 3dvolreg.

### 4.4 General procedure

The experiment consisted of 5 sessions: 1 familiarization session outside of the MRI scanner, 1 finger pattern localizer session inside the MRI scanner, and 3 decoded neurofeedback sessions inside the MRI scanner. Each fMRI session had a behavioral pre- and post-test. A custom-made force-sensitive keyboard was used inside the MRI scanner to control finger coordination patterns during neurofeedback. We used a within-subject control by providing 2 different types of neurofeedback on alternating trials: one based on index-middle finger coordination and one based on ring-little finger coordination.

### 4.5 Individual finger pressing task

An individual finger pressing task was performed during all fMRI scanning sessions (Fig 1B). This task was also performed for 15 minutes during the familiarization session. Each trial of the task consisted of 10 seconds of individual finger pressing followed by 2 seconds of feedback. Each trial was preceded by a 2 second cue for the target finger, and feedback was both preceded and followed by 1 second wait periods, leading to a 16 second trial duration. In the familiarization and localizer sessions, feedback was related to maintaining normal finger coordination patterns on the individual finger pressing task, while in the neurofeedback sessions, feedback was related to neural activity in sensorimotor cortex. Each fMRI session consisted of 8 runs of 20 trials each, lasting about an hour.

Participants were instructed to press the target finger (one of index, middle, ring, or little) while keeping a constant pressure on the 3 other (non-target) fingers. During the cue and pressing period, the real-time force of each finger was displayed as a grey disk (Fig 1B). For the target finger, a yellow target cylinder was visible indicating the target force range. Two types of targets were displayed in alternating order: low force (between 0.2N and 1.2N) and high force (between 2.0N and 3.0N). Participants were required to keep the grey finger disk in the target cylinder for 200ms before moving on to the next target (e.g. for high force targets, maintaining a force between 2.0N and 3.0N for 200ms). Participants had 10 seconds to hit as many targets as possible, indicated by a timer bar at the bottom of the screen. For each non-target finger, a cylinder was visible indicating a constant force range to maintain (0.2-1.5N). While fingers exerted a force in this range, the non-target cylinder turned green, and the grey disk was hidden by the cylinder; if fingers exerted more or less force than this force range, the cylinder turned red and the current force of the finger was visible as a grey disk, either above or below the cylinder depending on whether the force was less or greater than the desired force range.

Points were tallied during this task based on hitting targets with the target finger and maintaining constant force on the non-target fingers. Specifically, targets were assessed as ‘hits’ (+1 point, green score message: ‘+1’), ‘misses’ (0 points, red score message: ‘miss!’), or ‘non-target fingers off’ (0 points, red score message: ‘off!’). Hits were tallied if the target finger remained inside the yellow target cylinder for 200ms and all non-targets fingers remained inside the constant force cylinders. If the target finger entered and exited the target cylinder in less than 200ms, a miss was recorded. If the target was hit but any of the non-target fingers exited their constant force cylinder, then a ‘non-target finger off’ was recorded. Participants were encouraged to gain at least 15 points each trial. At the beginning of each trial, a grey score bar appeared at top of the screen, at the same width as the timer bar. As targets were hit, the score bar decreased in size, up to a score of 15. After 15 targets were hit, the score bar turned green and increased in size. After 10 seconds of pressing, a feedback thermometer appeared in the position of the target finger, related to the participant’s performance during that trial. If at least 15 targets were hit, the feedback thermometer cylinder turned green; if less than 15 targets were hit, the feedback thermometer was grey. Feedback was either related to the number of targets hit (familiarization and finger pattern localizer sessions) or the participant’s neural activity during pressing (neurofeedback sessions).

#### 4.5.1 Behavioral feedback

In the familiarization and finger pattern localizer sessions, the feedback thermometer was directly related to the number of targets hit. Initially, the maximum height of the thermometer was equivalent to 15 points. For each finger, the maximum score achieved by the participant in that session was used to recalibrate the maximum height of the thermometer. The numerical score was also shown below the feedback thermometer during the feedback period (in green for scores 15 or above or in grey for scores below 15). At the end of each run, the total score over all trials was shown at the bottom of the screen.

#### 4.5.2 Individual finger decoding

The fMRI data from the finger pattern localizer task was used to create a finger decoder to be used in the subsequent neurofeedback sessions. Standard preprocessing (rigid body motion correction, detrending, and z-scoring) was performed. The preprocessed fMRI data from the last 6 seconds of pressing was averaged over time for each trial to yield 160 time-averaged trials, 40 per finger. The hand area of the primary sensorimotor cortex was identified in each participant by first training a sparse multinomial logistic regression decoder (SMLR; PyMVPA) on the larger primary sensorimotor cortex (Brodmann areas 4A, 4P, 3A, and 3B identified using Freesurfer), and then selecting only those voxels within a 15mm radius of the center of the hand area. A final SMLR decoder was trained on only these voxels, and exported for use in neurofeedback. For decoded neurofeedback sessions, each run began with 20 seconds of baseline rest. At each TR, data was linearly detrended using the current time history and z-scored using the variance estimated from the initial baseline period.

#### 4.5.3 Neurofeedback

In neurofeedback sessions, participants were rewarded up to $30 per session for shifting the neural activity pattern of the middle finger towards that of the index, and for shifting the neural pattern of the ring finger towards the little finger. Specifically, the neurofeedback score is summarized in Equation 1, where *finger*_*reward*_ is the decoder output for the finger adjacent to the target finger that is being rewarded (middle: index; ring: little) and *finger*_*punish*_ is the decoder output for the finger adjacent to the target finger that is being punished (middle: ring; ring: middle). Time points used for decoding were identical to the finger pattern localizer task (i.e. the time-averaged fMRI data from the last 6 seconds of pressing).

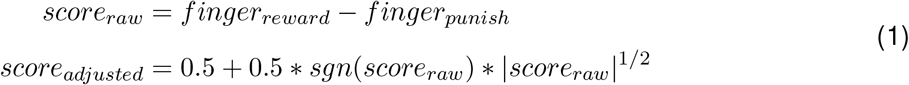

Participants were simply instructed to regulate their brain activity during pressing to maximize the height of the feedback thermometer (*score*_*adjusted*_), and that the neural activity signature would be different for each finger. If they achieved at least 15 points on the individual pressing task, they saw their neural activity feedback score as a green thermometer, indicating reward proportional to the height of the thermometer; if they did not achieve this minimum score, they saw their neural activity as a grey feedback thermometer and did not receive reward. Trials were blocked in groups of 5 repetitions of the same finger (i.e. 5 trials of middle finger presses followed by 5 trials of ring finger presses). The amount of money earned was displayed at the bottom of the screen at the end of each run.

#### 4.5.4 Post-hoc calculation of bias scores

For all statistical analyses and plots, bias scores were recalculated offline to ensure that they were not skewed by real-time processing. Variance for z-scoring was estimated using the baseline period at the beginning of each run, as in the neurofeedback sessions. Offline detrending was performed using GLM correction that accounted for the explicit finger presses during the neurofeedback sessions (Kopel et al., 2019). The same SMLR decoder and feedback calculation equation as in the neurofeedback sessions were then applied.

#### 4.5.5 Calculation of univariate activation

Univariate activations were calculated as percent signal change (PSC) for each finger in each run of each session. Because we were concerned about the impact of univariate activation on bias scores, the univariate region of interest (ROI) for each participant included only voxels that had a non-zero decoder weight for at least one finger. The mean activation from this ROI was extracted from the motion corrected data, without any z-scoring or detrending. A design matrix was constructed with a regressor for each finger as well as a constant and linear term for detrending estimation. Finger regressors were constructed as a boxcar during each 10-sec of pressing, convolved with a double gamma canonical hemodynamic response function. A least-squares fit between the model and univariate activation was then calculated with Python and Numpy (`numpy.linalg.lstsq`). PSC was estimated for each finger in each run as the model estimate for each finger divided by the model’s constant term estimate. PSC was calculated for each session by averaging the eight run estimates.

### 4.6 Behavioral assessment: rapid reaction time task

We assessed short-term plasticity associated with each neurofeedback session using a motor confusion task in which participants performed rapid button presses (700ms between presses) of pseudorandomly cued fingers (Fig 2C). Participants viewed the real-time force of each finger as a grey disk that moved up and down with each finger. Under each finger’s disk was a corresponding target grey disk. Every 700ms, one of the target disks would light up white, indicating the finger to be pressed. A selected finger was detected as the first finger to exceed a force of 1N. Participants were encouraged to make rapid presses of the correct finger through a point system. If an incorrect finger was selected, the target disk lit up red and a score of −1 point was tallied. If the correct finger was pressed with a reaction time greater than 450ms, the target disk lit up yellow and a score of +1 point was tallied. If the correct finger was pressed with a reaction time faster than 450ms, the target disk lit up green and a score of +2 points was tallied. At the end of each run (217 presses), the participant’s score was displayed. The press order in each run was randomized to ensure an equal number of each possible finger transition (18 transitions for each of the 12 possible finger transitions). Participants performed 6 runs of this task (approximately 15 minutes) immediately before and after each fMRI session in a room adjacent to the scanner.

### 4.7 Behavioral assessment: temporal order judgment task

Long-term changes in hand representation were assessed using a temporal order judgment task in which participants had to judge which of two adjacent fingers received a vibrotactile stimulus first (Fig 2D). This task was conducted during the familiarization session, as well as after the final neurofeedback session. Each run of the task targeted 2 adjacent fingers (index-middle, middle-ring, or ring-little). Participants rested their fingers on top of vibrotactile stimulators (Soberton E-12041808) which lay on the force-sensitive keyboard. The real-time applied force of the 2 targets fingers were displayed as grey disks. To begin each trial, participants were required to apply a constant force on the stimulators (between 0.2N and 0.5N, indicated by a blue cylinder) for 250ms. After this, the display went blank for 800ms. During this blank period, the two fingers were stimulated one after the other (20 ms duration, 200 Hz sinusoidal pulse), with possible inter-stimulus intervals (ISIs) of +400, −400, +250, −250, +180, −180, +120, −120, +70, −70, +30 or 30ms where, by convention, a positive ISI indicates the rightward digit was stimulated first. The onset of the first pulse was randomly jittered, starting between 150ms and 350ms after the beginning of the blank period. After the blank period, a target disk appeared in white for each finger, and the real-time force of each finger also re-appeared as a grey disk. Participants were instructed to select the finger which was first stimulated. Selections were detected as the first finger to exceed a force of 1N. The selected target lit up blue, and a blue asterisk appeared above the selected finger; however, no feedback was given as to whether the selection was correct. At this point, the two blue cylinders appeared again, and the next trial began once the participant kept the constant (between 0.2N and 0.5N) force on the stimulators for 250ms. At the end of each run (120 trials, 10 per ISI), summary accuracy over the run was displayed to participants. In each session of the temporal order judgment task, participants completed 9 runs total (3 for each finger pair) in pseudorandomized order, lasting about 45 minutes. Participants listened to pink noise over headphones to mask any auditory cues from the vibrotactile stimulators.

### 4.8 Finger coordination assessment

Coordination between the target finger and the adjacent (rewarded and punished) fingers during neurofeedback was assessed on a trial-by-trial basis. Each trial was subdivided into a number of presses equal to the number of high force targets passed. For the target finger, the force of each press was evaluated as the maximum force within +/−200ms of the high force target being passed, minus the minimum force within +/−200ms of the previous low force target being passed. Because participants were required to stay in each target for 200ms, this window ensured that the peak and trough forces of each press were typically captured. The timing of the maximum and minimum force in the target finger were then used to evaluate the corresponding pressing force of the adjacent fingers. Coordination was assessed as the mean pressing force of the adjacent finger (over all presses within the trial) divided by the mean pressing force of the target finger. Note that this coordination metric (evaluated as a ratio of forces) can be either positive or negative because the press timing is determined only by the peaks and troughs of the target finger presses, which may be out of phase with the peaks and troughs of the adjacent fingers (e.g. if participants are slightly lifting the adjacent finger during pressing of the target finger). For each participant, a heatmap of the coordination of the rewarded and punished fingers was generated using a bivariate kernel density estimate (Fig 4, lower panels, with Python and Seaborn’s kdeplot).

### 4.9 Statistical analyses

Paired t-tests were used to assess participants’ ability to bias the neural activity pattern associated with each finger relative to baseline, as well as to directly compare between fingers. A linear mixed effects model with participant as a random effect was used to assess the interaction between neurofeedback training and finger (ring or middle). For all bias score analyses, the mean bias score for each finger over all three neurofeedback sessions was compared to the mean values calculated for the pattern localizer session. Bias scores were recalculated offline, taking into account the explicit finger pressing task during neurofeedback, to ensure that they were not skewed.

Linear mixed effects models with participant as a random effect were used to determine the relationship between univariate activations and finger bias scores. A separate model was constructed for each finger (middle or ring). A general linear hypothesis test was used to calculate p-values for the slope of each model.

The relationship between baseline variability and participants’ ability to bias patterns was calculated using Spearman’s rho. Significance was established using 10,000 bootstrap correlations selecting either the ring or middle finger randomly for each participant, with the p-value calculated as the proportion of iterations with rho<0.

Paired t-tests were used to assess within-finger changes in finger preference at each neurofeedback session. Significance was assessed using Bonferroni adjusted alpha levels of 0.0125 per finger (0.05/4). Changes in preference were also compared directly between fingers with a linear mixed effects model (with participant as random effect) of the change in finger preference for each session, using a general linear hypothesis test of adjacent finger contrasts (index-middle, middle-ring, and ring-little). Changes in motor confusion between finger pairs at each neurofeedback session was assessed by creating a linear mixed effects model for each session with participant as a random effect. Tukey’s post-hoc test (*α* <0.05) was used to determine if there were statistically significant differences between finger pairs. A similar linear mixed effects model was created for assessing changes in tactile discrimination behavior due to all three neurofeedback sessions.

A linear model using coordination of each of the adjacent fingers was used to assess the effect of average coordination patterns on mean neurofeedback scores. A linear mixed effects model with participant as a random effect was used to assess the effect of trial-by-trial coordination patterns on trial-by-trial neurofeedback bias scores, again using the coordination of the two adjacent fingers as fixed effects. Finally, a general linear hypothesis test was used to calculate p-values for linear and linear mixed effects models.

#### Statistical software

For all analyses, linear mixed effects models were constructed with `lme` from `nlme` in R (Pinheiro et al., 2018). Linear models without mixed effects were constructed with `lm` from base R. Interaction effects were assessed with anova from base R. Post-hoc tests for main effects were conducted using `glht` from `multcomp` in R (Hothorn et al., 2008). Basic statistics (paired t-tests, Spearman’s rho) were calculated with Python’s `scipy.stats`.

### 4.10 Subjective assessments and questionnaires

A neurofeedback strategy questionnaire was administered after each neurofeedback session to determine the strategies used by participants for each finger. At the end of the last neurofeedback session, participants were asked what they thought the feedback signal for each finger was related to, and were finally given a forced-choice test to guess which non-target finger they thought the feedback thermometer was related to (for each target finger).

## Notes

### Competing Interest Statement

The authors have declared no competing interest.

### Summary of Updates

Added further statistical comparison between fingers; added sample size justification

